# Prioritization of Deleterious Mutations Improves Genomic Prediction and Increases the Rate of Genetic Gain in Common Bean (*Phaseolus vulgaris L.*), a Simulation Study

**DOI:** 10.1101/2025.05.05.652208

**Authors:** H. Cordoba-Novoa, V. Hoyos-Villegas

## Abstract

The study of mutations is fundamental to understanding evolution, domestication, and genetics. Characterizing mutations has the potential to accelerate breeding programs through selection and purging of deleterious mutations (DelMut). Here, we investigated how predicting DelMut in breeding populations can improve genomic prediction (GP) and inform strategies to increase the rate of genetic gain. DelMut were annotated in three independent common bean populations using a previously developed random forest (RF) model incorporating phylogenetic and protein information. Deleterious scores from the RF model were mostly around 0.25, with the top 1% (*highly* DelMut) of variants scoring between 0.78 – 0.82 among populations. All populations showed variation in the number of *highly* DelMut per line (max. 13 – 197) and in genetic load. We assessed the impact of incorporating a priori information for variant prioritization and weighting based on predicted deleteriousness in GP models for yield and flowering time. Stochastic simulations were conducted to evaluate how different mating schemes based variable numbers of DelMut per parent affect genetic gain. Variants with higher predicted scores had significantly different effect distributions compared to random or lower-scored markers. Yield predictions were 4.47–12.3% more accurate when markers were weighted by effect and deleterious score; no consistent improvement was observed for flowering time. Simulated breeding cycles showed that selecting parents with fewer *highly* DelMut consistently increases the rate of genetic gain. These results highlight the potential of DelMut information for variant prioritization and the optimization of common bean breeding programs. The approaches we developed can be assessed in other species to improve the efficacy of crop improvement.

**Key messages:**

- Predicted deleterious mutations have different distributions of effects based on population composition.
- Variant prioritization and differential weighing of markers based on effects and deleterious scores can improve the prediction of yield.
- Favoring mating schemes between parents with fewer highly deleterious mutations can increase the rate of genetic gain.

## Introduction

The study of mutations plays a paramount role in understanding the evolution and adaptation of species, as well as genetic and phenotypic variation (Charlesworth et al., 1993; Gossmann et al., 2012; Piganeau and Eyre-Walker, 2003). Depending on their effect, mutations can be neutral, advantageous or deleterious. The distribution of the effects of mutations helps explain the proportion of mutations that can be beneficial or detrimental depending on environmental forces (Eyre-Walker and Keightley, 2007; Krasovec et al., 2016). Nonsynonymous mutations (both advantageous or deleterious) are under strong selective pressures. Strongly advantageous mutations are usually rare (Eyre-Walker, 2006) and highly deleterious mutations (DelMut) are purged (Crow, 1970). However, mildly recessive DelMut with subtle effects can escape purifying selection and are accumulated, known as the genomic genetic load (Glémin, 2003).

Multiple factors affect the accumulation of mildly DelMut. In crop species, domestication and selection bottlenecks contribute to the increase of inbreeding in populations, which in turn reduces the effective population size and effectiveness of purifying selection, leading to the accumulation of weakly DelMut (Charlesworth and Charlesworth, 1999; Kono et al., 2016). The accumulation patterns and potential role of DelMut in phenotypic variation are topics of particular interest in crop improvement. Studies on the characterization of DelMut in diverse populations have been conducted in several species with varying accumulation patterns depending on the mating system and breeding methods (Dwivedi et al., 2023). In self-pollinated species such as common bean and soybean, breeding has reduced the number of DelMut, but mildly DelMut are fixed and accumulated (Cordoba-Novoa et al., 2025; Kim et al., 2021). The accumulation of DelMut can limit selection and hinder the genetic gains in breeding programs (Moyers et al., 2018; Zhu et al., 2022).

The identification and characterization of DelMut in crop species raises the question of how genetic gain in breeding programs can be further accelerated (Johnsson et al., 2019; Wallace et al., 2018). The targeted removal of DelMut using new genomic techniques (NGT) such as genome editing is one of the avenues for crop improvement (Gao, 2021; Glaus et al., 2025; Johnsson et al., 2019). Other approaches aim to enhance the prediction ability (PA) of genomic selection (GS) models. Some studies have considered the use of predicted DelMut to subset markers for GS or their inclusion as fixed effects in the model (Valluru et al., 2019; Wu et al., 2023). Methods that modify the relative importance of markers in the model through the inclusion of posterior variances or the modification of genomic relationship matrices (GRM) have also been considered (Edwards et al., 2016; Long et al., 2023; Yang et al., 2017). The different approaches have shown varying levels of efficacy depending on the trait, the populations, and the environments.

In breeding programs, decisions and final results are simultaneously influenced by various parameters. Stochastic simulations have been adopted as tools to simulate breeding scenarios and probable outcomes for optimizing breeding programs (Covarrubias-Pazaran et al., 2022; Hassanpour et al., 2023). Such simulations have the advantages of recreating entire populations with genotypic and phenotypic data at the individual level, which provides precise predictions of the consequences of proposed changes at different stages, such as crossing, evaluation, and selection (Liu et al., 2019; Vieira et al., 2025). Stochastic simulations have been employed for the simulation and optimization of animal (Gorjanc et al., 2018; Johnsson et al., 2019), and plant breeding programs, including rice (Fritsche-Neto et al., 2024), sweet corn (Peixoto et al., 2024), soybean (Silva et al., 2021), and common bean (Chiaravallotti et al., 2024; Lin et al., 2023) considering the adoption of GP frameworks.

As a predominantly self-pollinating species, common bean may accumulate DelMut with varying effects. As mildly DelMut can accumulate in different genomic regions, the purging of DelMut from breeding populations and elite material has the potential to increase the effectiveness of breeding programs. However, no studies on the potential practical applications of the inclusion of DelMut in plant breeding pipelines have been conducted. Thus, we investigated the potential impact of using information derived from the prediction of DelMut for optimizing breeding programs. Our objective was to predict DelMut in different common bean breeding populations and evaluate how genomic prediction (GP) could benefit from variant prioritization in the model. Additionally, we hypothesize that selecting parents and designing mating schemes based on a priori information about the presence of DelMut could accelerate the rate of genetic gain over time in breeding programs.

## Materials and Methods

### Phenotypic and genotypic data

Three independent common bean breeding populations with publicly available phenotypic and genotypic data were used in this study. The populations have different genetic backgrounds, breeding histories, and genotypic datasets, and have been evaluated in different environments. The black bean MAGIC population was developed at McGill University, Canada, with eight parents from the Middle American Diversity (MDP) panel (Moghaddam et al., 2016). Founders were selected to maximize allelic diversity and crossed in a multi-funnel scheme (half-diallel). A total of 18 families and 532 recombinant inbred lines (RIL) were advanced. Field experiments were conducted in two locations in Canada in 2024 (Cordoba-Novoa et al., 2025). For the present study, the evaluations from the Sainte Anne de Bellevue, QC (SAB) location were used based on data quality and the experimental design. Best Linear Unbiased Estimators (BLUE) were calculated fitting a mixed-effect model in the *lme4* R package (Bates et al., 2015), and a Complete randomized block design (CRBD)Click or tap here to enter text., and means were corrected for further analysis. Flowering was evaluated as the number of days from planting until 50% of the plot plants had at least 50% of their flowers open (DTF). Yield was recorded as Kg/Ha based on the plot weight. The MAGIC RILs were skim-sequenced using 150 paired-end PCR-free libraries in the Illumina platform and imputed using a Practical Haplotype Graph (PHG) built from Nanopore and Illumina Reads from the eight founders (Bradbury et al., 2022; Cordoba-Novoa et al., 2025). A total of 1.5 million SNPs were available for the present study.

The second population was the Vivero Equipo Frijol (VEF). VEF is an elite Andean population that is part of the bean breeding program of the International Center of Tropical Agriculture (CIAT). Details on the population and phenotypic evaluation can be found in Keller et al. (2020). For the present study, we used the VEF phenotypic data for 346 genotypes evaluated in Darien (Valle del Cauca, Colombia) under the middle phosphorous level (DAR16C_mdP location). The location was selected among the locations with the greatest number of evaluated genotypes and with similar prediction abilities in the environment (DAR16C) where VEF was evaluated. BLUE-corrected means for DTF and yield (Kg/Ha) were available and directly used for our analysis. As previously reported, the VEF population was genotyped using Genotype-by-sequencing (GBS) with the *ApeKI* restriction enzyme and Illumina sequencing. After quality control, 5,820 SNP markers were available.

The third population was the Cooperative Dry Bean Nursery (CDBN), a multi-environment trial (MET) grown for over 70 years in the US and Canada. MacQueen et al. (2020) reported the data analysis strategies for four decades of the CDBN evaluation. For yield, we used phenotypic data from 318 genotypes evaluated at the MSTI (Sidney, Montana, US) location, and for DTF, we used data from 226 genotypes evaluated at the WYPO (Powell, Wyoming, US) location. These locations had the highest number of observations (datapoints) for each trait. Due to the unbalanced nature of the data, Best Linear Unbiased Predictors (BLUP) were calculated across years. MacQueen et al. (2020) genotyped the CDBN accessions by re-analyzing raw sequencing from previous reports and sequencing some genotypes with a dual-enzyme (*MseI* and *TaqI*) GBS approach. After variant calling and QC, a total of 1.2 million SNPs were available.

### Population structure and LD analyses

As mentioned above, the populations used were previously reported and characterized to a certain extent. To facilitate the analysis of the results, we evaluated population structure using principal component analysis (PCA) implemented in Tassel v5 (Bradbury et al., 2007) and pairwise linkage disequilibrium (LD) in GAPIT v3 (Wang and Zhang, 2021). LD was plotted using the R package LDheatmap (Shin et al., 2006). For population structure and LD evaluation, the genotypic data sets of the MAGIC population and CDBN were reduced to a similar number of markers by setting a minimum distance of 10 Kbp between SNPs while maintaining the LD patterns.

### Prediction of putatively deleterious mutations

To identify putatively DelMut, two Random Forest (RF) models for common bean were reported by Cordoba-Novoa et al. (2025), one trained based on the Middle American and another on the Andean Common bean genome reference. The Middle American RF model was used for the MAGIC population, whereas the Andean RF model was used for the VEF and the CDBN. The model choice depends on the genetic background and the reference genome (UI111 – Middle American or G19833 – Andean) used for variant calling in each population. Briefly, for the model implementation, Sorting Intolerant From Tolerant For Genomes (SIFT4G; Ng and Henikoff, 2003) scores and unified representation (UniRep) embeddings (Alley et al., 2019) are obtained for each genotypic dataset and later loaded in the RF models to calculate a deleterious score. The higher the value from zero to one, the more deleterious an allelic change is predicted to be. The SNPs in the top 1% of the deleterious scores within each population were classified as *highly* deleterious (*highly* DelMut).

To obtain a genome-wide estimation of the deleterious burden of each genotype in each population, the homozygous (Hom), heterozygous (Het), and total genetic load per genotype was estimated as the sum of all the predicted scores for a line according to the allelic state (i.e. if the derived allele is not present (or is the reference, the score is not added) following a similar approach as Wu et al. (2023).

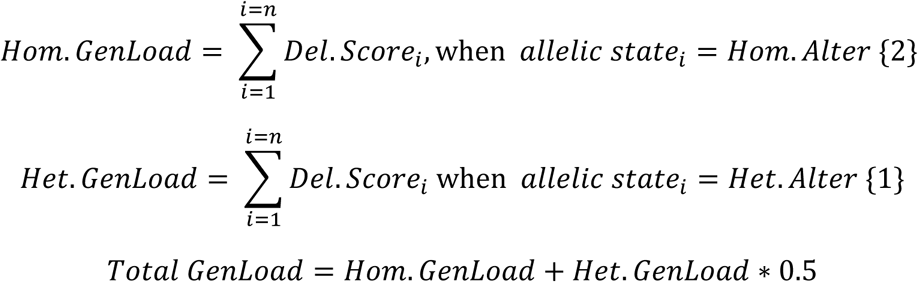

### Estimation of effect sizes

Following the approach proposed by Valluru et al. (2019) the effect size of the SNPs was estimated using a RR-BLUP model in the R-package rrBLUP v4.6 (Endelman, 2011).

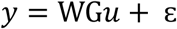

Where ***y*** is the vector of BLUEs or BLUPs for yield or DTF, **W** is the matrix relating individuals to observations, **G** is the genotype matrix coded as {-1,0,1} under an additive model, and ***u* ∼** *N*(0, I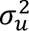) is the vector of marker effects. The marker effects can be written as *û* = *Z′*(*ZZ′* + *λI*)^−1^*y*, where Z=WG and λ is the ridge regression parameter defined as the ratio between residual and marker variances (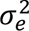/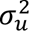).

For each population, the distributions of the effects of SNPs with and without score were compared. Since the number of SNPs with scores is lower, random subsets of equal size from SNPs with no scores were defined. The minor allele frequency (MAF) was verified among subsets to avoid bias. SNPs Scored for DelMuts were further subset, and effects re-calculated to compare the effect distributions between the SNPs with the top 30% DelMut score and the remaining 70% based on weight distribution and percentiles. Differences between distributions were evaluated using the Kolmogorov–Smirnov (Ks) test in R.

### Genomic prediction

To evaluate the potential of incorporating information from DelMut into genomic prediction (GP), a Genomic Best Linear Unbiased Prediction (GBLUP) and a Bayesian Ridge Regression (BRR, also known as Gaussian prior) model were used. The following GBLUP model was implemented in ASReml (Butler et al., 2023).

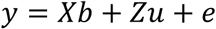

Where *y* is the vector of phenotypic BLUEs or BLUPs, **X** is a design matrix relating the fixed effects to each genotype, *b* is the vector of fixed effects, **Z** is a design allocating the records of genetic values, *u* is the vector of additive genetic effects for a genotype, and *e* is the vector of random normal errors. In the model, *var*(*u*) = 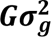 where **G** is the genomic relationship matrix (GRM) and 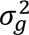 is the genetic variance for the model. The GRM was constructed using the additive method from VanRaden (2008) in the R package *AGHmatrix* (Amadeu et al., 2023) as follows:

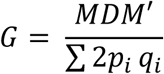

Where **M** is the centered genotype matrix coded as 0, 1 and 2 for the homozygous reference allele, heterozygous, and homozygous alternative allele, respectively. **D** is the identity matrix, and *p* and *q* are the allele frequencies.

The following BRR model was implemented in the BGLR package (Pérez and De Los Campos, 2014).

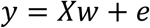

Where **X** is the design matrix, and *e* is the independent and identical distributed Gaussian error. **w** is the vector of marker effects (model parameters) where **w** ∼ *N*(0, *λ*^−1^I). In the model, *y* is the vector of phenotypic values with 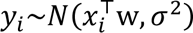. In the BRR model, the number of Markov Chain Monte–Carlo iterations per model was 12,000 with the first 2,000 as burn-in without partition of training and testing datasets.

To evaluate model performance, each population was randomly split into 70% and 30% for the training and testing datasets. The predictions were compared based on the results from a 10-fold cross-validation with 50 iterations for GBLUP and 10 for BRR. The average prediction ability (PA) was determined by Pearson’s correlation between the predicted and the observed phenotypic values. Mean PAs were compared using Student’s t-test.

### Inclusion of deleterious mutations information in Genomic Prediction

For the inclusion of DelMut information in the GBLUP model, new GRMs were constructed weighing the SNPs based on their effect and predicted DelMut scores. The SNP weights are the absolute value of the estimated effect (from the rrBLUP model) times the predicted score:

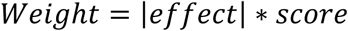

SNPs with the same or similar estimated effect will be considered differently in the model depending on how deleterious (expressed by the score) the allelic change is predicted to be. SNPs with no DelMut scores were kept unweighted in the model. The GRMs were weighed according to Liu et al. (2020) for the VanRaden method used.

Unlike GBLUP, the BRR model starts from the genotype matrix rather than a GRM. To inform the model about DelMut, the marker dataset was split into scored and unscored SNPs and defined as two different random effects in a two-level list in the model, following a multi-kernel approach (Pérez and De Los Campos, 2014). To avoid potential size bias, in each iteration the SNP subsets were randomly sampled and kept to the same number of markers.

### Genomic relationship matrix dimension reduction

For the VEF population, no further filtering or reduction was applied to the SNP dataset. Based on the original GP study for VEF that showed that the marker number (higher than 1K markers) did not affect the model PA, and to give enough room to accommodate SNPs with deleterious scores, subsets of 3K SNPs were used for the iterations (Keller et al., 2020). Due to the high number of SNPs and to alleviate computational burden, the MAGIC and CDBN genotypic datasets were reduced using the *--indep-pairwise* tool in Plink v1.9 (Chang et al., 2015) with a window size of 50Kb, step of 1 and a r^2^ threshold of 0.3, similar to previous reports for common bean (Keller et al., 2020). After reducing the MAGIC and CDBN marker datasets, SNPs with scores were re-incorporated into the baseline dataset if they had been deleted during the pruning step. In each iteration of these two populations, we used subsets of 7K randomly selected markers, maintaining an equal number of SNPs with and without deleterious scores (weighted and unweighted, respectively).

### Simulation of the genetic gain

We evaluated how incorporating information on DelMut could improve selection and increase the rate of genetic gain over time. As proof of concept, we simulated a hypothetical basic breeding scenario using a pedigree breeding strategy with artificial selection for a quantitative trait in each generation (represented in Lin et al., 2023). Here, the initial crosses of the first breeding cycle are carried out between parents solely selected based on the number of *highly* DelMut. Three independent crossing schemes (closed programs) were compared. First, genotypes with a high number of *highly* DelMut were crossed between them **High × High** (**H × H**); second, parents with a low number of *highly* DelMut were crossed **Low × Low** (**L × L**); and third, parents with a high number of *highly* DelMut were crossed with parents with a low number of *highly* DelMut **High × Low** (**H × L**). An additional scheme of random crosses was also simulated. Within each proposed scenario, the top (higher number of DelMut) and the bottom (lower number of DelMut) 20 parents were used for the initial random crosses. For subsequent breeding cycles (second onwards), parents are selected from the advanced lines of the previous cycle based on the phenotype (top simulated parents are included in the next cycle). After cycle 0 was complete, no further selection based on DelMut content was made among the set of parents that made up cycle 1 onwards. In the simulations, 10 breeding cycles from initial crosses to advance yield trials (AYT) were simulated with 50 iterations in AlphaSimR (Chris Gaynor et al., 2021). Real (not simulated) genotypic and phenotypic data collected for each population (VEF, MAGIC, CDBN) were used for the simulations. Average genetic gain was calculated per cycle as the difference between the mean genetic value of the varieties and mean genetic value of the parents in each cycle. Cumulative genetic gain was calculated in each cycle by adding the previous cycle’s shift in genotypic value. To keep comparisons stable among populations, broad-sense heritability (H^2^) in AlphaSimR was set as 0.25 for yield and 0.45 for flowering based on previous reports for common bean (Delfini et al., 2021; Kamfwa et al., 2015; Keller et al., 2020; Raggi et al., 2019). The genetic map reported by Diaz et al. (2020) was used for simulations.

## Results

### Population structure and LD

The three populations, VEF, MAGIC, and CDBN, have different breeding histories, genetic backgrounds and were genotyped using various methods, which makes direct comparisons challenging. However, different population structures and LD patterns were observed within each population. In the PCA, VEF had a main cluster with a few lines on the far right and explained variation of 22.4% between PC1 and PC2 (Figure 1A). The MAGIC population had two main subpopulations, mainly based on the PC1, that explained 9.5% of the variation (Figure 1B). The CDBN had three clear subpopulations tightly clustered with PC1 and PC2, explaining up to 52.4% of the variation (Figure 1C).

**Figure 1.**
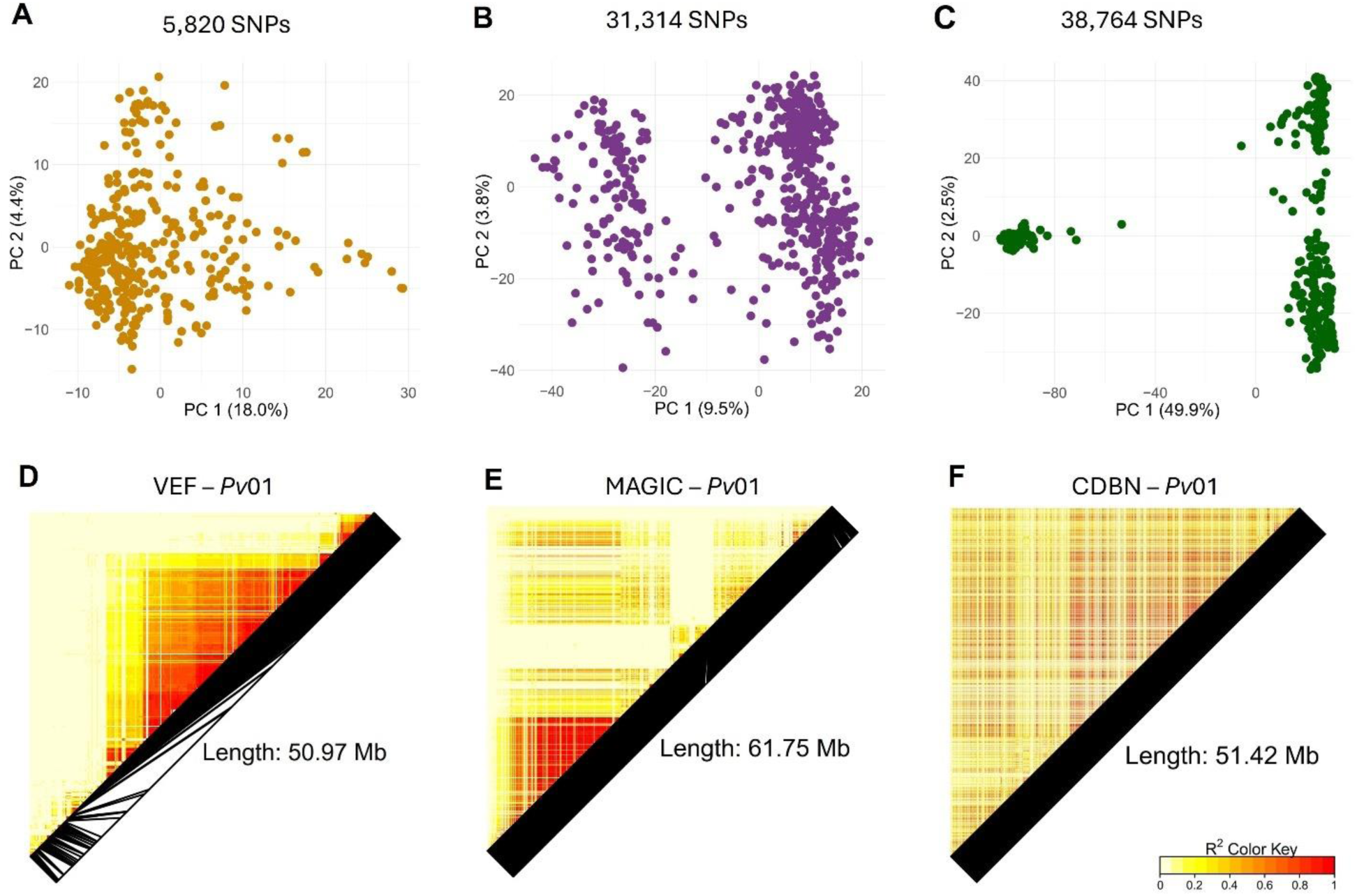
Principal component analysis (PCA) for population structure in Vivero Equipo Frijol (VEF; A), Black bean MAGIC population (B), and Cooperative Dry Bean Nursery (CDBN; C). Linkage disequilibrium (LD) heatmaps for chromosome 01 (*Pv*01) in VEF (D), MAGIC (E), and CDBN (F). The SNP dataset is independent for each population. LD patterns for the other chromosomes in Figure S1-3.

Despite differences in marker density, similar chromosome lengths were observed in the LD analysis of each population. Different LD patterns were identified in each population (Figure S1-3). Figure 1 D-F shows the LD heatmap for *Pv*01 in each population as an example of the varying patterns observed in each population. For instance, a big cluster of markers in high LD (r^2^ > 0.8) was observed in the middle of Pv01 for VEF, while a smaller cluster, still in high LD, was identified at the beginning of the chromosome for MAGIC. In CDBN, no clear and large LD blocks were observed.

### Phenotypes

Descriptions of the phenotypic data can be found in the original reports for each population (Cordoba-Novoa, 2025; Keller et al., 2020; MacQueen et al., 2020). Briefly, in VEF, yield ranged from 457.5 to 1,815 Kg/Ha with a mean of 1,072.6 Kg/Ha (CV = 22.9%). Flowering ranged between 35.4 to 40.2 days and a mean of 38.2 days (CV = 2.5%). In the MAGIC RILs, yield ranged between 544.7 – 3,177.5 Kg/Ha, with a mean of 1,886.2 Kg/Ha (CV = 20.1%), and DTF from 41.5 to 50.5 days and a mean of 45.7 days (CV = 3.9%). In the CDBN dataset, yield in the MTSI location had higher values compared to VEF and MAGIC, ranging from 2,063.3 to 3,832.8 Kg/Ha and a mean of 3,065 Kg/Ha (CV = 9.9%). In the WYPO location, DTF varied from 54.3 to 62.2 days, with a mean of 58.4 days (CV = 2.3%).

### Prediction of deleterious mutations in populations

The number of annotated SNPs with deleterious scores varied in each population depending on the data availability. For the VEF population, 1,072 SNPs had a predicted score, 4,753 in the MAGIC population, and 14,740 in the CDBN dataset. Scores varied from zero to one, with most of the values around 0.25 in all three populations (Figure 2A-C). Due to a higher marker density, in the CDBN there was a wider score distribution. Based on the top 1%, the threshold for *highly* DelMut was 0.82 for VEF and CDBN, and 0.78 for MAGIC. In the three datasets, the number of *highly* DelMut was between 1.18 – 1.34% of the SNPs with a predicted score. For the total genetic load, the three populations showed normal distributions with values between 250 – 350 for VEF, 200 – 600 for MAGIC, and 4400 – 5200 for CDBN (Figure 2D-F; Table S1-3). Interestingly, in the distribution of the CDBN, only a few lines within the Middle American group (n = 15) had total loads between 4800 and 5000, with two peaks in the distribution (Figure 2F; Table S3). The homozygous genetic load calculated only with the *highly* DelMut showed the same distribution patterns for the three populations (Figure S4).

**Figure 2.**
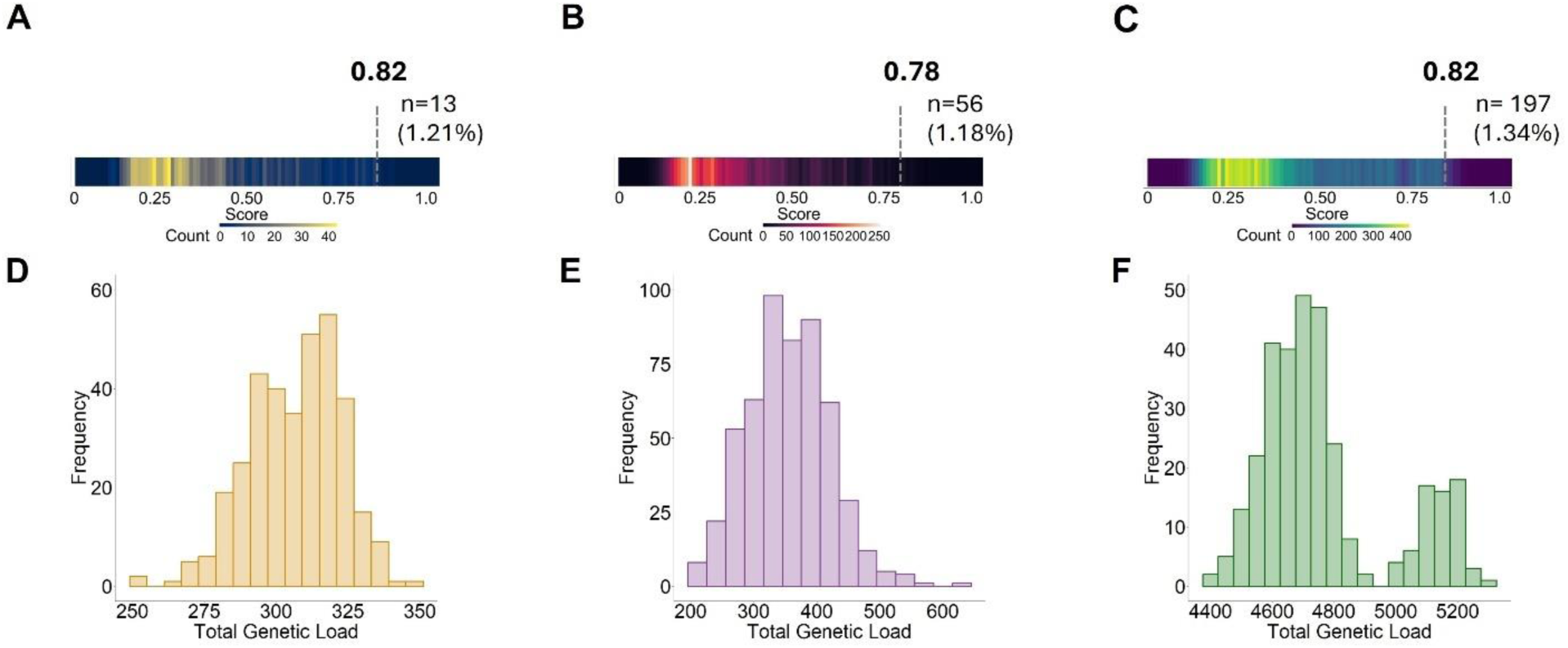
Prediction of deleterious mutations in three common bean breeding populations, Vivero Equipo Frijol (VEF; A, D), Black bean MAGIC population (B, E), and Cooperative Dry Bean Nursery (CDBN; C, F). Distribution of deleterious scores (A-C) and distribution of total genetic load (D-F). The SNP dataset is independent for each population.

Scored SNPs accumulated in 800 genes for the VEF population (Table S4), 7,639 for MAGIC (Table S5), and 4,943 in CDBN (Table S6). *Highly* DelMut accumulated in 12 genes in VEF, 37 genes in MAGIC, and 143 genes in CDBN with 1-3 mutations per gene (Table S7).

### Estimation of effect sizes

The distributions of the absolute values of SNP effects showed differences between scored and unscored SNPs for yield and DTF. For yield, the effects of unscored SNPs were more evenly distributed compared to the scored SNPs. The effects for scored SNPs were mainly concentrated around zero, with a highly leptokurtic distribution and higher peaks (Figure 3). Significant differences (*p* < 0.001) between the distributions were observed in the MAGIC and CBDN populations, with marginal differences in VEF (*p* < 0.05). When the scored SNPs were split based on the 30% percentile score threshold (0.39 for VEF, 0.38 for MAGIC, 0.5 for CDBN), a similar pattern was observed (Figure S5). Markers with a higher score (predicted to be more deleterious) had a near zero peak compared to SNPs with lower score values. For DTF, the same behavior for the distribution of scored and unscored SNPs was observed in the three populations (Figure 4). When comparing scored SNPs for the CDBN dataset, opposite to the trend, SNPs with scores < 0.5 had lower effects with a higher peak compared to SNPs with scores > 0.5 (Figure S6 E-F).

**Figure 3.**
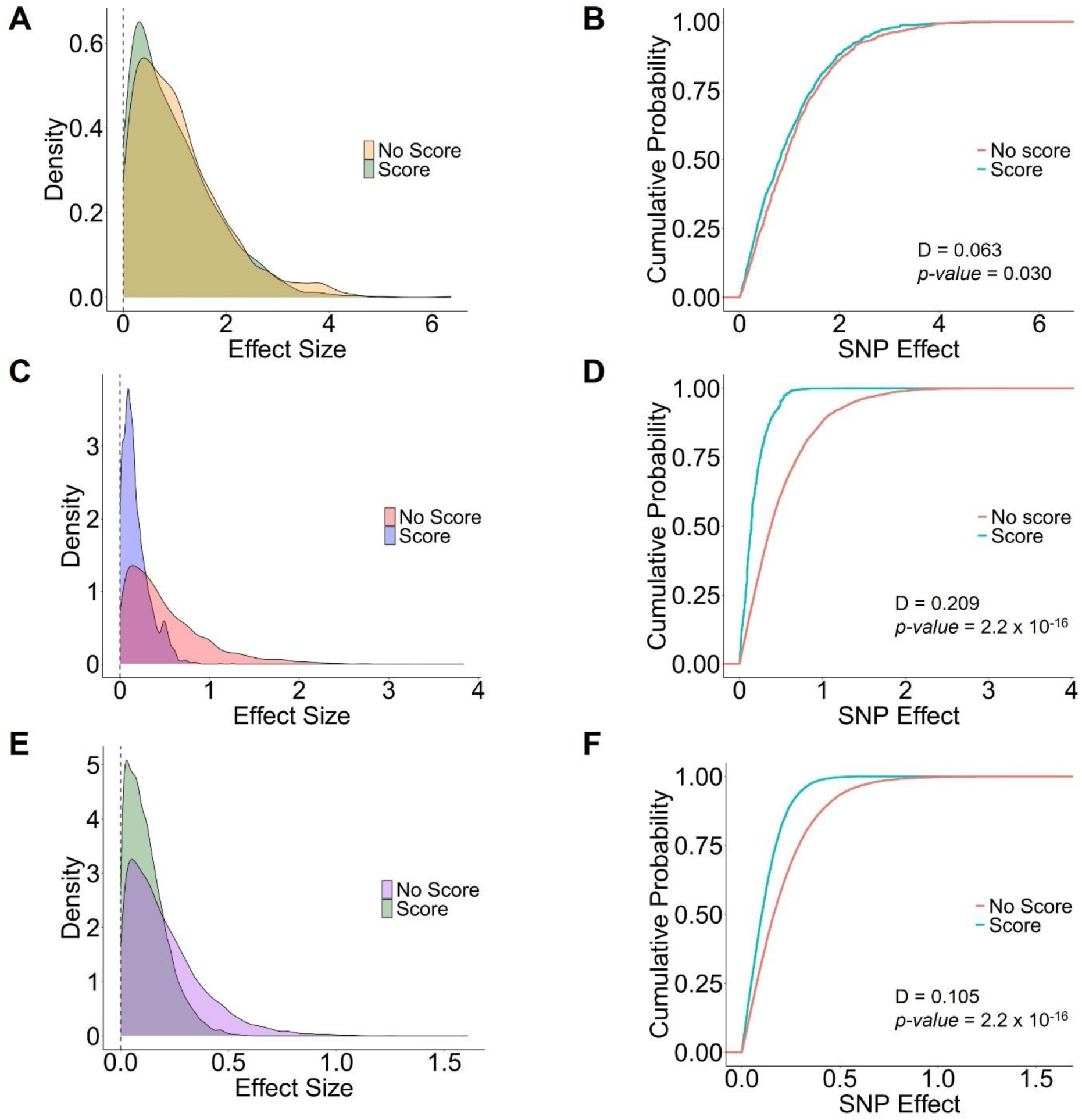
Distribution of the absolute value of the estimated effect of scored and unscored SNPs for yield in Vivero Equipo Frijol (VEF; A), Black bean MAGIC population (C), and Cooperative Dry Bean Nursery (CDBN; E). Cumulative probability for the distribution of the effects (B-F). D and p-value correspond to the Kolmogorov–Smirnov (Ks) test for distributions. The SNP dataset is independent for each population.

**Figure 4.**
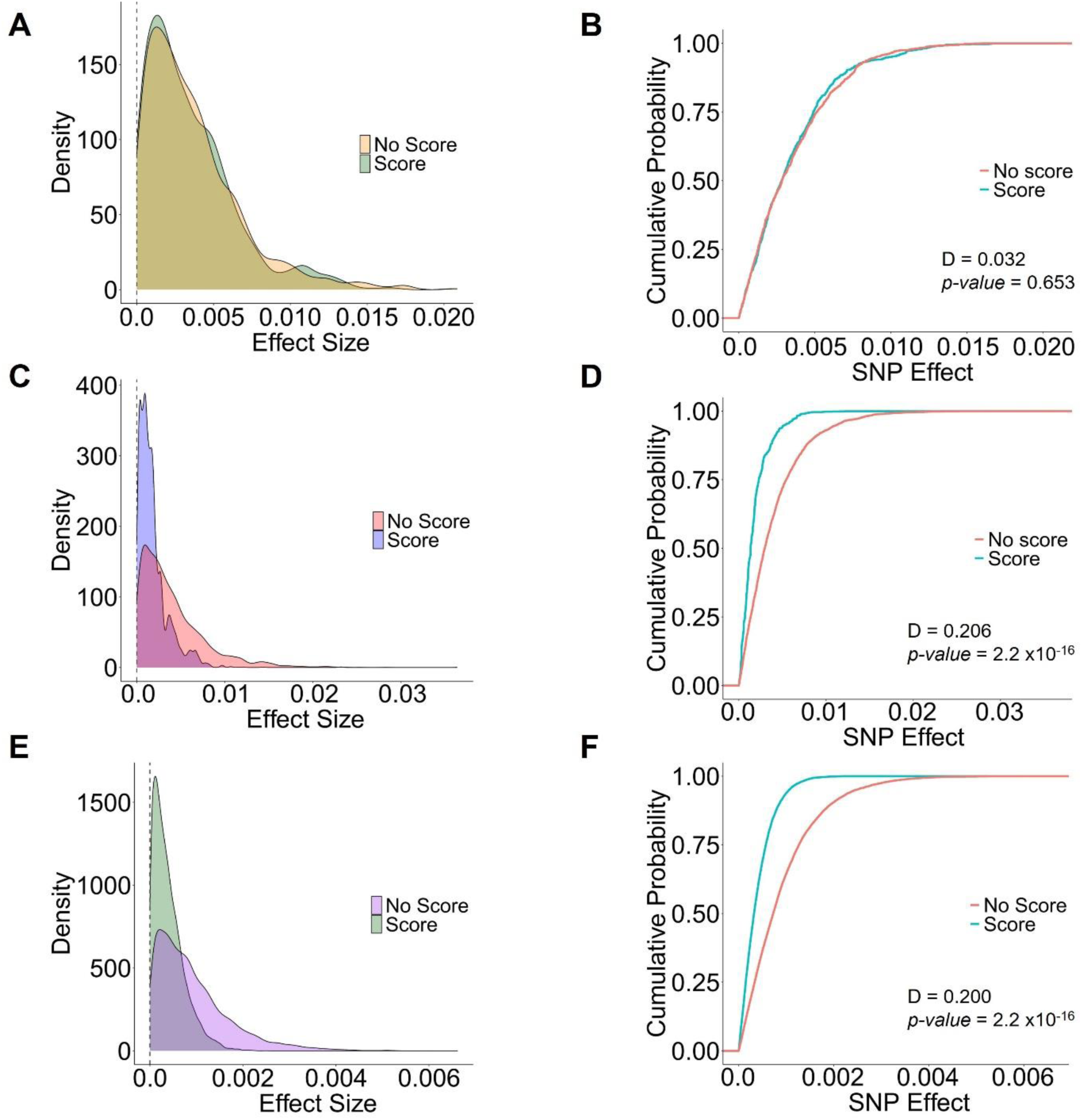
Distribution of the absolute value of the estimated effect of scored and unscored SNPs for days to flowering (DTF) in Vivero Equipo Frijol (VEF; A), Black bean MAGIC population (C), and Cooperative Dry Bean Nursery (CDBN; E). Cumulative probability for the distribution of the effects (B, D, F). D and p-value correspond to the Kolmogorov–Smirnov (Ks) test for distributions. The SNP dataset is independent for each population.

### Genomic prediction

We investigated how incorporating previous information on DelMut could improve the prediction ability of genomic prediction models. For yield, the unweighted GBLUP and simple BRR had similar results with higher prediction abilities in the CDBN dataset (0.50 - 0.51), followed by VEF (0.38 - 0.40), and MAGIC (0.25). When incorporating the information from DelMut (weights based on effects and deleterious score for GBLUP, and multi-kernel for BRR), the prediction ability increased for all three datasets using GBLUP, and for VEF and CDBN using BRR. The weighted GBLUP model resulted in a 12.3% increase in the PA in the VEF population (0.52), 5.45% (0.31) in MAGIC and 4.47% (0.55) in CDBN. The multi-kernel BRR increased the PA by 6.5% in VEF and 3.0% in CDBN (Figure 5A). Significant differences were only detected for GBLUP.

**Figure 5.**
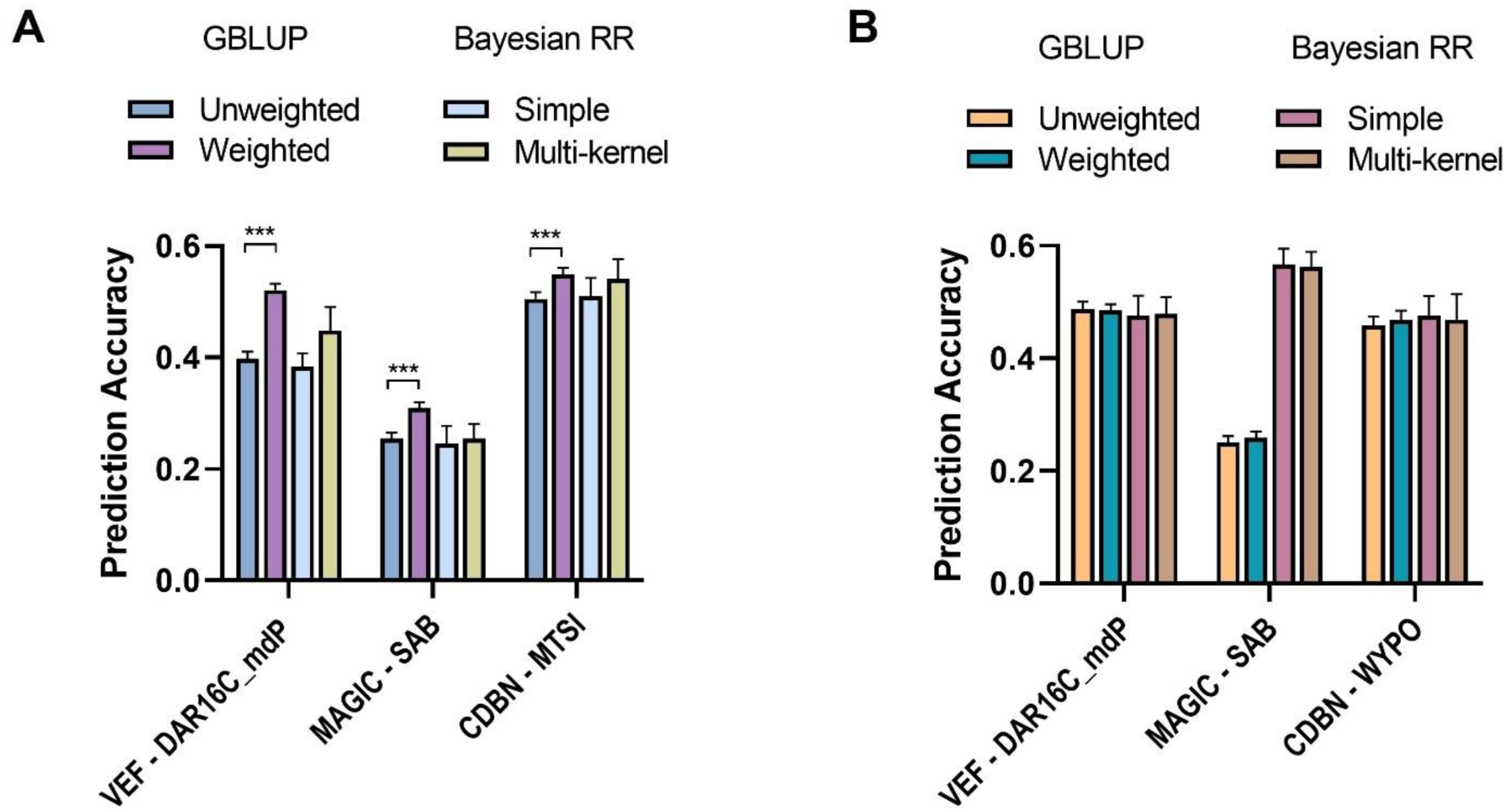
Prediction accuracy for yield (A) and days to flowering (B) using weighted and unweighted GBLUP, and simple and multi-kernel BRR models for the Vivero Equipo Frijol (VEF), Black bean MAGIC, and Cooperative Dry Bean Nursery (CDBN) populations. *** indicates significant differences (*p* < 0.0001) according to student’s t-test.

For flowering (DTF), the unweighted GBLUP had PA of 0.49 for VEF, 0.46 for CDBN, and 0.25 for MAGIC. Similar PAs were obtained for VEF and CDBN using the simple BRR model. However, a higher PA for DTF in MAGIC was obtained using the simple BRR model (0.57). The incorporation of weights in GBLUP marginally improved the PA of the models for DTF in MAGIC (0.88% higher) and CDBN (0.93% higher). The multi-kernel BRR model had similar results to the GBLUP models with a marginal increase of 0.23% in the PA in VEF (Figure 5B).

### Simulation of the genetic gain

Genotypes from each population were selected as potential parents to simulate new *closed* breeding cycles with a pedigree method. Parents were selected solely based on the number of *highly* DelMut alleles. In the VEF population, parents with a “Low” number had seven to eight mutations and “High” had 12-13 (Table S8). In the MAGIC population, selected parents in the “Low” group had 0-3 mutations, and those in the “High” group had 16-19 (Table S9). From the CDBN population, “Low” DelMut genotypes had 16-19 mutations, and *“*High*”* genotypes had between 31-33 (Table S10). Random simulated crosses behaved similarly to the H × L scheme, the results were excluded to facilitate visualization and comparison of the defined mating systems based on DelMut.

The genetic gain for yield decreased every cycle in each population. In the first breeding cycle of the derived population from VEF, the genetic gain was 1.37 Kg/Ha in the H × H scheme, 2.17 Kg/Ha in the H × L scheme, and 2.22 Kg/Ha in the L × L. In the second cycle, L × L showed a higher genetic gain (1.63 Kg/Ha) compared to the other two strategies (0.39 – 0.67 Kg/Ha). The increase in genetic gain was close to zero after the fifth cycle for H × L and H × H crossing schemes, whereas small increases were observed for the L × L strategy until the seventh cycle (Figure 6A). In the cumulative genetic gain, higher increases were observed for the crosses made from parents with a low number of *highly* DelMut (L × L) with a final gain of 5.85 Kg/Ha (0.54%) followed by the H × L crosses with 4.16 Kg/Ha (0.39%), and finally the H × H crosses with 2.43 Kg/Ha (0.23%).

**Figure 6.**
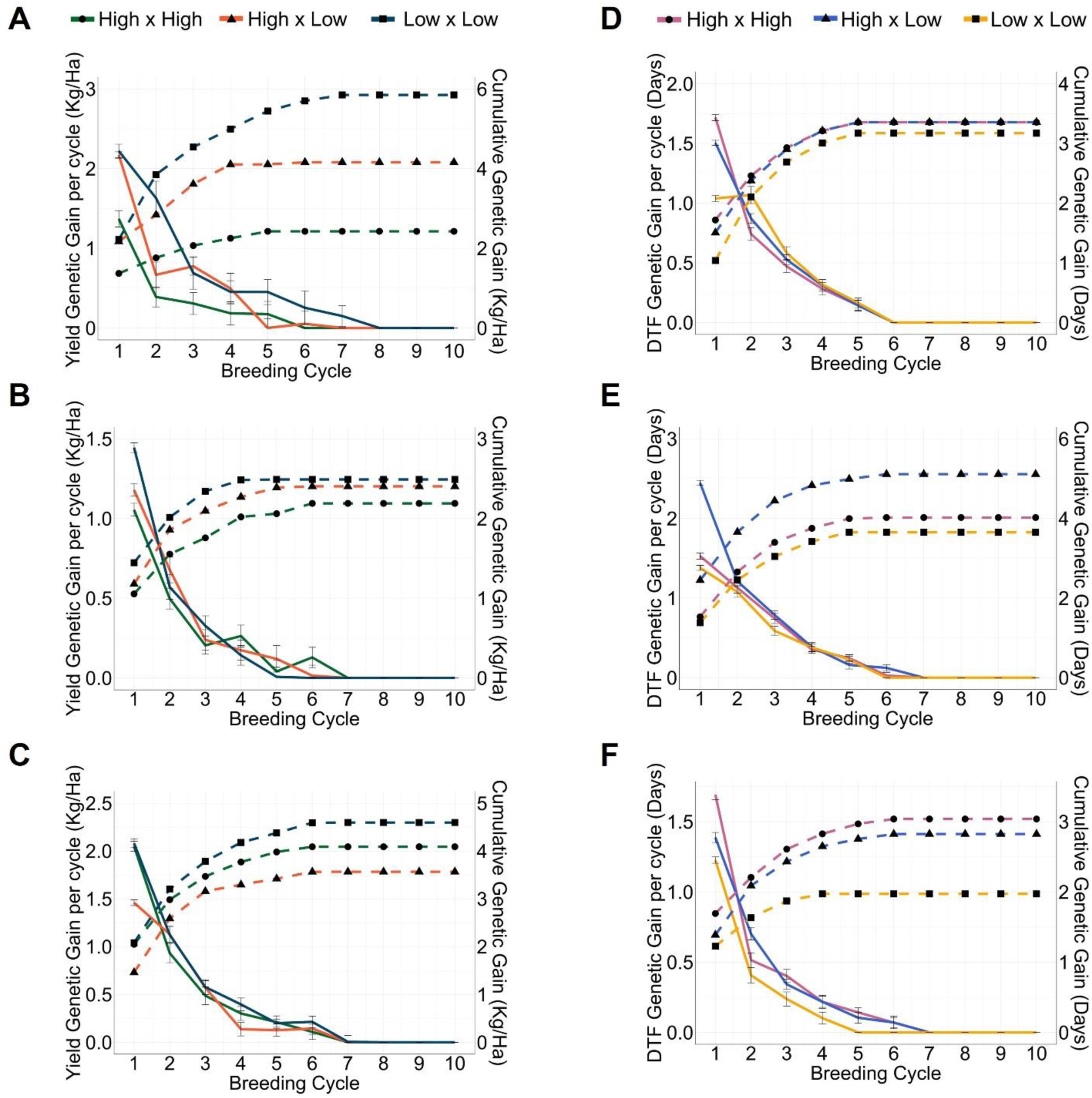
Simulation of genetic gain per cycle and cumulative genetic gain for yield (A-C) and days to flowering (D-F) for the Vivero Equipo Frijol (VEF; A, D), Black bean MAGIC (B, E), and Cooperative Dry Bean Nursery (CDBN; C, F) populations. High × High, High × Low, and Low × Low indicate the crossing schemes based on the number of *highly* deleterious mutations in each population.

Similar trends in the reduction of genetic gain per cycle and the increase of cumulative genetic gain were observed for all the selection strategies in the MAGIC and CDBN populations for yield. In the MAGIC population, lower genetic gains were predicted (Figure 6B). In the first cycle, the yield increase for the L × L scheme was 1.44 Kg/Ha, 1.17 Kg/Ha for H × L, and 1.05 Kg/Ha in H × H. By the 10^th^ cycle, the cumulative genetic gain was lower in H × H (2.18 Kg/Ha; 0.11%) compared to both L × L (2.49 Kg/Ha; 0.13%) and H × L (2.40 Kg/Ha; 0.13%).

In the simulation from the CDBN genotypes, the genetic gain in the first cycle was around 2.06 Kg/Ha for the L × L and the H × H crossing schemes, followed by the H × L with 1.46 Kg/Ha. The final cumulative genetic gain across crossing strategies was lower than VEF (but higher than in MAGIC). As in VEF and MAGIC, the cumulative genetic gain was highest in the L × L scheme, reaching 4.59 Kg/Ha (0.15%). However, the H × H derived crosses had a final genetic gain of 4.09 Kg/Ha (0.13%) followed by the H × L scheme with 3.57 Kh/Ha (0.12%; Figure 6C).

Low genetic gains for flowering were observed in the simulated cycles from the VEF and CDBN populations. In the VEF simulations, H × H and H × L had higher genetic gains in the first cycle (1.72 and 1.5 days, respectively) compared to L × L (1.04 days). By the sixth cycle all schemes no longer showed a genetic gain per cycle (Figure 6D). Final cumulative genetic gains were similar for H × H and H × L (3.35 days; 8.5%) and slightly lower for L × L (3.17 days; 8.0%).

For the MAGIC and CDBN simulations, clearer differences among crossing schemes were observed. In MAGIC, higher genetic gains for flowering compared to VEF and CDBN were observed. In the first cycle, the H × L crosses gained 2.45 days, H × H 1.53 days, and L × L 1.38 days. This trend between crossing schemes was maintained in the cumulative genetic gain, with a final increase of 5.11 days (10.7%) in H × L, 4.01 days (8.5%) in H × H, and 3.65 days (7.7%) in L × L. For the CDBN simulation, the L × L strategy had lower genetic gains in DTF compared to the other two and reached zero at the fourth cycle. The H × H and H × L schemes registered gains until the sixth cycle (Figure 6F). The final cumulative gain in DTF was higher in H × H (3.04 days; 5.1%) and H × L (2.82 days; 4.8%) than in the L × L cycles (1.97 days; 3.3%).

## Discussion

### Effect of deleterious mutations

We investigated how the prediction of DelMut can potentially inform selection decisions and accelerate breeding programs. We evaluated three publicly available common bean breeding populations with diverse genetic and breeding histories. Methods for the prediction of DelMut based on conservation constraints have been previously developed (Davydov et al., 2010; Ng and Henikoff, 2003). The adopted approach has the advantage of combining both phylogenetic information and the likely impact of amino acid substitutions, providing a continuous classification of deleteriousness for variant prioritization (Cordoba-Novoa et al., 2025; Long et al., 2023; Ramstein and Buckler, 2022). The observed scores had an expected distribution where most of the annotated SNPs have low values around 0.25. Mutations with a low score are likely to be tolerated, while mutations with a high score are predicted to be *highly* deleterious (Cordoba-Novoa et al., 2025; Long et al., 2023).

When considered individually, the effects of DelMut may seem inappreciable. It is the aggregated effect of multiple putatively DelMut with subtle individual effects that may have an impact on the overall plant fitness (Felsenstein, 1974; Kono et al., 2019). The distribution of the fitness effects is a research topic itself and explains the proportion of mutations that may be deleterious, neutral or advantageous within populations (Eyre-Walker and Keightley, 2007). Mutation accumulation experiments and other studies have shown that the effects of mildly DelMut generally follow a leptokurtic gamma distribution with a shape parameter (*α*) lower than one (Eyre-Walker et al., 2006; Keightley, 1996). A highly leptokurtic gamma distribution was observed for the scored SNPs (and above certain threshold) in yield and flowering (Figure 3 and 4). This indicates that most of the mutations are mildly deleterious, and a few of them have large (positive or negative) fitness effects in the long right tail of the distribution (Keightley, 1996). These results are consistent with the distribution of the predicted scores (Figure 2A-C) and the observed distributions within scored SNPs (Figure S5-6). Additionally, Ohta and Kimura (1971) suggested that mutations are not equally deleterious but range from near-neutral to slightly deleterious to very deleterious alleles.

Leptokurtic distributions with low shape parameters are characterized by the clustering of values at the minimum (Low effect; Böndel et al., 2022; Piganeau and Eyre-Walker, 2003). In the biological context of DelMut, large-effect mutations that can be lethal or highly detrimental to plant fitness are under strong purifying selection and eventually eliminated from the population. However, small-effect mutations can be tolerated and accumulated at a higher rate. Domestication and breeding reduce effective population sizes, increase inbreeding, and reduce the effectiveness of purifying selection, which leads to the accumulation of tolerated mildly and a few *highly* DelMut (Grossen and Ramakrishnan, 2024; Kono et al., 2016). Mutations with high effects and scores are predicted to have more significant impacts on plant fitness compared to other variants and may be more informative for selection approaches.

### Genomic prediction

The prediction of phenotypes from genotypic data using trained models on related populations has been of extensive interest in plant and animal breeding. Multiple factors affect the PA of models. Strategies such as variant selection and differential marker weighing have been proposed to improve genomic selection efforts.

There are multiple factors affecting the prediction accuracy (PA) of GP, such as the training and testing populations, model selection, and variant selection (Alemu et al., 2024; VanRaden et al., 2017). GBLUP is a widely used model that has been demonstrated to generally be superior in predicting different traits (Wang et al., 2018). On the other hand, with a Bayesian framework, BRR directly models marker effects, could be more flexible, and has been used for GP in common bean (de los Campos et al., 2013; Keller et al., 2020). GBLUP assumes that all markers contribute equally to the total genetic variance, modeling breeding values through a genomic relationship matrix. BRR, in contrast, assumes normally distributed marker effects with equal variance, estimating them directly in a Bayesian framework. Here, we consider the fact that small- and large-effect DelMut can accumulate at different rates in breeding populations and how conservation and predicted deleteriousness can assist variant prioritization to optimize GP. In our results, differentially weighting SNPs based on their effects and predicted deleteriousness scores in GBLUP, as well as partitioning scored and unscored SNPs as separate random effects in the BRR model, improved the prediction accuracy (PA) of GBLUP for yield across all three evaluated populations (Figure 5A). VanRaden et al. (2017) and Xavier (2019) highlight the importance of variant selection, pruning, and filtering to enhance genomic prediction. Causal variants and markers closer to them or in strong linkage are expected to explain higher proportions of variance compared to random markers (Meuwissen et al., 2024). Very conservative filtering methods may result in losses in PA, while an excessive number of markers increase computational burdens.

Previous reports have shown that weighted GP approaches can improve prediction models in animal and plant species with varying success (Fang et al., 2017; Long et al., 2023; Meuwissen et al., 2024; Zhang et al., 2024). Yield is a highly complex trait controlled by multiple small-effect genes and their interactions. One characteristic of the scored SNPs from the RF model is that they are in coding regions (Cordoba-Novoa et al., 2025; Ramstein and Buckler, 2022). Prioritizing putatively DelMut with individual small effects could help the model capture a substantial and biologically relevant *cumulative* effect (de los Campos et al., 2013; Goddard et al., 2010). Another plausible explanation is that prioritized SNPs (scored and weighted) can be in a strong linkage disequilibrium (LD) with unknown causal variants, contributing to a better capture of the genetic variation and prediction (Vilhjálmsson and Nordborg, 2012). Kono et al. (2019) showed in barley that top progeny selected based on an RR-BLUP model had fewer DelMut relative to other lines. This suggests that DelMut affect the observed phenotypes and GP models indirectly account for putatively DelMut. In this regard, in other self-pollinating species such as soybean, elite lines have been shown to have a lower number of DelMut (Kim et al., 2021). Negative correlations between DelMut (and genetic loads) and yield have also been reported for common bean (Cordoba-Novoa et al., 2025), maize (Yang et al., 2017), and potato hybrid populations (Wu et al., 2023).

The VEF population was previously evaluated in multiple environments and traits using a genomic prediction framework. The PAs we obtained for yield and flowering with either GBLUP or BRR follow the previous study (Keller et al., 2020). Keller et al. (2020) found that the inclusion of significant QTL signals as fixed effects did not improve the PA of the model. Instead of accounting for fixed effects, our approach directly modifies how SNPs are accounted for in the model based on the effect and/or deleterious score, showing consistent PA improvements in VEF and other populations for yield.

Predictions for days to flowering (DTF) did not benefit from incorporating DelMut information (weights or multi-kernel partition) into the model. Flowering time is a less complex trait than yield, with a larger proportion of phenotypic variance attributed to genetic factors (higher broad-sense heritability). As a less complex trait, flowering time is controlled by fewer, bigger-effect genes compared to yield. A higher number of small-effect genes involved in the multiple subprocesses affecting yield also leads to a higher accumulation of DelMut affecting the trait (more genes accumulate more DelMut related to the trait). This may explain why the prediction of yield is more responsive to the inclusion of DelMut information in the GP models.

In contrast to yield, significant QTL and genes identified through linkage mapping and GWAS have been reported for DTF (Ates et al., 2018; Nascimento et al., 2018; Raggi et al., 2019). Given that these QTLs account for a significant percentage of the phenotypic variation, additive genetic effects may already be effectively captured in the model, with no apparent advantages from SNPs with minor effects, such as the scored ones. When examining the effects of marker selection for GP in radiata pine (*Pinus radiata*) and shining gum (*Eucalyptus nitens*), Klápště et al. (2020) also noted that low heritability traits benefit more from model refinement. In simulation studies, Morgante et al. (2018) demonstrated that the prediction of traits governed by a greater number of QTLs (more complex) are more susceptible to parameter tuning compared to traits of lower complexity.

### Simulation of the genetic gain

Stochastic simulations support the optimization of breeding programs under varying parameters. Mating system designs influence the efficiency of breeding cycles and the rate of genetic gain over time. In our simulated closed system, the selection of parents based on the number of *highly* DelMut influenced the expected genetic gain over time. For yield, higher per-cycle and cumulative genetic gains were consistently observed in all the populations for breeding schemes where the initial parents had a low number of *highly* DelMut (**L × L**). Since the genetic load of plants does not have evident phenotypic effects and DelMut are not routinely considered in GP schemes, the genome-wide prediction and later consideration of DelMut in breeding materials can inform selection decisions, particularly in the choice of parents. The incorporation of the knowledge about DelMut is of especial importance in animal breeding. For instance, Sonesson et al. (2003) and Raoul et al. (2018) stress the value of selecting against carriers and designing mating systems based on the incorporation or exclusion of these individuals. Carriers are individuals known to carry recessive mutations associated with diseases that could express later in the breeding process.

Johnsson et al. (2019) simulated how the selection against carriers and genome editing of DelMut increase animal average fitness. They observed that the most effective strategy depends on the DelMut level of dominance (codominant or recessive). Selection against carries is more effective for recessive mutations, whereas genome editing has the potential for both. Different gene action models have been proposed for DelMut, including additive, incomplete dominance, and codominance (Robinson et al., 2023; Sun et al., 2023). In an additive action model, the small effects of the predicted DelMut are independent and add up linearly, which may contribute to the observed improvements in GP and the final simulated genetic gain in each crossing scheme. In other species such as maize, incomplete dominance of DelMut can also contribute to trait variation and heterosis (Yang et al., 2017). Additional research on other gene action effects of DelMut may open new avenues on how effects are modeled and then incorporated in breeding programs.

In our cumulative genetic gain of DTF, breeding initiated with parents with a low number of *highly* DelMut (**L × L**) had a lower gain. Short flowering time has been positively correlated with yield in previous studies (Kamfwa et al., 2015; Moghaddam et al., 2016). It is possible that when selecting against parents with a high number of *highly* DelMut, unknown mutations in genes involved in flowering are indirectly excluded, particularly in the first breeding cycle. For instance, the repair of a domestication DelMut in a floral regulator induced early fruit yield and more compact plants in tomato (Glaus et al., 2025). After the first cycle, parents are selected based on simulated phenotypes (top ones), which, based on the relationship between yield and flowering, can influence a lower flowering time in the cumulative genetic gain. Based on our results, the selection and design of mating (crossing) schemes between parents with a low number of predicted DelMut (**L × L**) has the potential to increase the genetic gain in yield and reduce flowering time.

The accumulation of DelMut can limit the rate of genetic gain in breeding programs (Moyers et al., 2018). Domestication and improvement increase inbreeding and reduce the effective population size, which in turns reduces the effectiveness of natural selection and leads to the accumulation of mildly and slightly DelMut (Dussex et al., 2023; Renaut and Rieseberg, 2015). Depending on the breeding purpose and the target environment and conditions, the effects of DelMut can be associated with their effect on the overall plant fitness. In a simplified manner, fitness can be defined as the ability of an organism to survive and reproduce in a specific environment (Orr, 2009). The level of fitness of an individual is related to the proportion of the next generation represented by its offspring, and depends on the contextual framework. In a breeding context, for instance, plants able to survive biotic and abiotic stress or yield more in different environments are expected to have a higher fitness. That same ability to produce more can be affected by the genetic load of the individual or the population if analyzed as a whole (Dwivedi et al., 2023).

Different responses to prediction and simulation were observed between populations. As mentioned above, characteristics proper to each population limit direct comparisons and restrict results within each one. In a practical setting, each breeding program can evaluate the presence and inclusion of DelMut in the context of the already-implemented genotyping and selection approaches. Our results show consistent patterns of how the information on DelMut can benefit both GP and the design of crossing blocks. The MAGIC population had a lower PA and simulated genetic gains in yield compared to VEF and CDBN. Multiparent populations are designed to accumulate a greater number of recombination events, and the progeny lines are mosaics with contributions of all parents (Gardner et al., 2016; Wang et al., 2022). These advantages make MAGIC populations particularly useful for gene mapping with a higher resolution, trait introgression, and marker-assisted selection. As more recombination events are accumulated, the linkage between alleles can be broken and allelic diversity increased. While this is desirable for eliminating DelMut in LD with beneficial alleles, this may also break the linkage between two or more positive alleles in coupling phase. In fact, Tourrette et al. (2019) and Taagen et al. (2022) showed that increased recombination, either in pericentromeric or chromosome-wide regions, can be detrimental for genetic gain and GP accuracies when it breaks up QTL in coupling phase. An increased recombination is particularly beneficial when positive QTL are in repulsion phase or targeted recombination approaches are adopted (Ru and Bernardo, 2019). These may partly explain why the MAGIC population had lower PAs and simulated genetic gains, especially for yield where more QTL are expected to contribute to the trait variation (more complex). Additionally, Keller et al. (2020) observed lower PA in a different MAGIC population compared to the same VEF population used here.

As proof of concept, the adopted approach has some limitations. In breeding programs, after each breeding cycle, new parents are integrated into the pipeline to preserve genetic diversity and continue increasing the genetic gain linearly. In our simulations, the genetic gain per cycle decreases after each cycle as alleles are fixed and the inbreeding increases. For the same reason, the cumulative genetic gain reaches a plateau value with no further increases. Here, we demonstrated *in silico* how the selection against DelMut has the potential to make breeding programs more efficient. Future simulations and empirical studies should evaluate simulations with the re-introduction of new germplasm similarly selected based on the number of DelMut. As a multifactor process, breeding programs are complex. Simulations that include changes in other parameters and evaluate the expected response of parents selected based on DelMut would also be valuable.

## Conclusions

We evaluated whether predicting deleterious mutations in common bean breeding populations can enhance genomic prediction (GP) and increase the rate of genetic gain for yield and flowering time. Variants with predicted scores and with higher values had a highly leptokurtic gamma distribution, different from that of random and lower-scored markers. Incorporating variant effects and deleterious scores as weights in the GRM of a GBLUP model or in a multi-kernel approach for a BRR model improved yield prediction by 3.0 – 12.3%, while marginal improvements were observed for DTF (0.23 – 0.93%). The selection of parents with fewer *highly* DelMut has the potential of increasing the genetic gain in yield and flowering. For instance, in the VEF population, crosses between genotypes with seven to eight *highly* DelMut had a higher cumulative genetic gain for yield (5.85 Kg/Ha) compared to crosses with 12-13 *highly* DelMut (2.43 Kg/Ha). For flowering time, crossing schemes with fewer *highly* DelMut also showed more favorable DTF changes. These findings demonstrate the potential of DelMut information to support breeding program optimization through GP and simulation-based approaches.

## Acknowledgements

We thank the Pulse Breeding and Genetics group members for their constructive feedback. We thank Isabella Chiaravalotti for the constructive discussions and guidance in the genomic prediction experiments, and Dr. Alice McQueen for facilitating the genotypic datasets for the CDBN population.

## Data availability statement

Code and materials used in this study can be found in the McGill University Pulse Breeding and Genetics GitHub page (https://github.com/McGillHaricots/peas-andlove)

